# Objective QC for diffusion MRI data: artefact detection using normative modelling

**DOI:** 10.1101/2023.06.30.546837

**Authors:** Ramona Cirstian, Natalie J. Forde, Jesper L.R. Andersson, Stamatios N. Sotiropoulos, Christian F. Beckmann, Andre F. Marquand

## Abstract

Diffusion MRI is a neuroimaging modality used to evaluate brain structure at a microscopic level and can be exploited to map white matter fibre bundles and microstructure in the brain. One common issue is the presence of artefacts, such as acquisition artefacts, physiological artefacts, distortions or image processing-related artefacts. These may lead to problems with other downstream processes and can bias subsequent analyses. In this work we use normative modelling to create a semi-automated pipeline for detecting diffusion imaging artefacts and errors by modelling 24 white matter imaging derived phenotypes from the UK Biobank dataset. The considered features comprised 4 microstructural features (from models with different complexity such as fractional anisotropy and mean diffusivity from a diffusion tensor model and parameters from neurite orientation, dispersion and density models), each within six pre-selected white matter tracts of various sizes and geometrical complexity (corpus callosum, bilateral corticospinal tract and uncinate fasciculus and fornix). Our method was compared to two traditional quality control approaches: a visual quality control protocol performed on 500 subjects and quantitative quality control using metrics derived from image pre-processing. The normative modelling framework proves to be comprehensive and efficient in detecting diffusion imaging artefacts arising from various sources (such as susceptibility induced distortions or motion), as well as outliers resulting from inaccurate processing (such as erroneous spatial registrations). This is an important contribution by virtue of this methods’ ability to identify the two problem sources (i) image artefacts and (ii) processing errors, which subsequently allows for a better understanding of our data and informs on inclusion/exclusion criteria of participants.

## 1. Introduction

Diffusion MRI (dMRI) is a neuroimaging modality frequently used to study the configuration of white matter in the brain. Diffusion refers to the molecular mobility of water molecules in biological tissue which can be measured in terms of its anisotropy levels. Due to the organization of white matter in bundles of myelinated axonal fibres, anisotropy can be exploited by dMRI to map the microscopic details of tissue architecture [1]. Although dMRI is a great tool for investigating in vivo structural organisation in the human brain, it is not without challenges. The resolution of a dMRI scan is typically lower than a regular T1-weighted anatomical scan as well as being more predisposed to the presence of artefacts. One of the main reasons artefacts arise is due to the increased sensitivity to off-resonance fields of the echo-planar imaging technique used for data acquisition. Likewise, the dynamic nature of collecting multiple volumes during a dMRI scan makes it susceptible to subject movement [2].

Several biophysical models can be applied to dMRI data to estimate tissue microstructure. Here we will briefly mention two of the most used models. The diffusion tensor imaging (DTI) model allows a full characterization of molecular mobility variation in space according to direction. Two of the most widely used DTI parameters are fractional anisotropy (FA) and mean diffusivity (MD) [3]. Neurite orientation dispersion and density imaging (NODDI) is another popular dMRI model. NODDI produces neurite density and orientation dispersion estimates which constitute more specific markers of brain tissue microstructure than standard indices than DTI [4].

DMRI data are affected by a slew of different artefacts [5], possibly more so than any other MRI modality. The scans are typically acquired using echo planar imaging (EPI) which means that they have a very low bandwidth along the phase-encode (PE) direction, of the order of 10-30 Hz/pixel. That means that they are very sensitive to, even very small, off-resonance fields resulting in geometric distortions. The dominating sources of off-resonance fields are i) Susceptibility, where the object (head) itself disturbs the main magnetic field by virtue of differences in magnetic susceptibility and ii) Eddy currents, where a field is generated by currents in conductors within the bore, currents that are in turn caused by switching of the diffusion gradients. The former, caused by the object itself, is mainly localised near the sinuses and ear-canals, and remains as a first approximation constant across all dMRI volumes. In contrast the second, caused by the diffusion gradient, affects the whole brain and is different for each volume. The field from either source causes geometric distortions (displacement of signal along the PE-direction) of several mm.

Because of the relatively long duration of a dMRI data set (comprising tens to hundreds of volumes) it is also affected by subject movement. The effects of this include “gross movement”, where the brain appears at a different place within the FOV in different volumes. That effect can be corrected by rigid-body registration. But, uniquely to diffusion MRI, movement can also cause full or partial loss of signal in individual slices (or groups of slices if Simultaneous Multi Slice (SMS) is used). The loss of signal is irreversible and correction methods are aimed at detecting it and minimising its effect on subsequent analysis.

The presence of artefacts in the dMRI scans may lead to problems in downstream processing and can bias subsequent analyses. For this reason, the artefacts should be corrected or removed in case correction is not possible. It is not uncommon to use a visual assessment protocol for dMRI datasets [6, 7, 8, 9]. This involves the visual inspection of each participant’s image either in terms of their full diffusion image and/or derived images such as FA and residual maps. Quality control protocols vary greatly and may include different degrees of rigour and complexity. In some cases, the artefacts are labelled by severity as well as type while other times a binary category is used (artefactual vs non artefactual). The labelling process requires a great deal of attention, expertise, and time, and it is ultimately subjective in nature due to the high variation in artefact appearance. For this reason, manual inspection of large datasets is not feasible and increases the risk of either discarding valuable data or including artefactual scans into a study.

Many automated quality control (QC) tools are part of a processing pipeline, which are often used for detecting and correcting artefacts in larger datasets. Most of the time these pipelines base their algorithm on the exclusion of artefactual data prior to further processing. The removal of data can be made at different levels of the image processing. For example, RESTORE [10] is a method for improving tensor estimation on a voxel-by-voxel basis in the presence of artefactual data points. The algorithm detects artefactual voxels by computing an initial tensor model using non-linear least squares and evaluating the residuals. PATCH [11] is a tool for the detection and correction of motion artefacts at the slice or patch level. It uses regional and more global (slice-wise) information to detect artefacts which improves the algorithms robustness and sensitivity. At the volume level, DTIprep [12] can detect and correct artefacts based on entropy estimation from all volumes. The volumes which reduce the entropy most are removed until the z-score for removal is below a set threshold. A common issue with this type of approach is the removal of too much data, which in turn may lead to a poorly estimated model or simply data loss.

Alternatively, there are pipelines which are based on estimating a desired outcome (e.g., average signal of the slice). For example, FSL EDDY [13,14] is a tool which retrospectively estimates various artefact types (eddy currents, susceptibility, movement, and slice dropout) by finding the distortion fields that achieve the best alignment of individual volumes to a model free prediction of what each volume should look like. Within the same framework, EDDY QC [15] is a tool for generating qualitative single subject and group reports, summarizing several objective QC parameters which are acquired based on the FSL EDDY output.

At the subject level, YTTRIUM [16] is a dMRI QC method which employs two QC metrics: 1) the skeleton-averaged (using TBSS) diffusion parameters values such as FA, MD and others in conjunction with 2) an estimate of the structural similarity between each subjects’ diffusion parameter map and the cohort average of that parameter (in MNI space). Using these two metrics, k-means is applied to estimate of the distance between each point (representing one subject) and the center of the cluster. A threshold of the k-means estimates separates the outliers who are affected by artefacts and/or errors from the non-affected subjects [16].

Recently, deep learning-based methods for quality control of neuroimaging data have gained popularity. In the field of diffusion imaging, a few notable works emerge. The QC Automator [17] is a method based on convolutional neural networks for automated classification of dMRI volumes and uses transfer learning for the classification of axial and sagittal slices. Furthermore, 3D-QCNet [18] is a more recent algorithm which improves upon the QC Automator principle by creating a deep learning pipeline that can detect dMRI artefacts three-dimensionally without requiring manually labelled data.

In summary, there are extensive dMRI image processing pipelines designed to minimize the effect of artefacts and correct the images after acquisition. Nevertheless, some challenges remain. After the pre-processing stage (e.g., denoising, deringing, susceptibility and motion correction), the data is further processed by applying different models, spatial registration and other steps where subsequent issues can also arise. Because of the many types of artefacts which can affect diffusion data, the detection of errors within the images is either time consuming and subjective, in the case of manually labelled scans, or may miss artefacts in the case of automated pipelines (e.g., classifiers). Many of these artefacts may be quite rare, which makes it difficult to obtain a sufficiently large, labelled dataset for training automated QC methods. Another issue with the existing pipelines arises from the many correction steps that are applied during the processing (registration to standard space). While some artefacts may be ‘corrected’, some are incompletely corrected or missed. Furthermore, approaches based on examining only the data may miss artefacts that occur during downstream processing, for example, spatial normalisation errors, which can occur more often in diffusion data than in other modalities due to the high precision required for brain structure localization (e.g., white matter tracts).

In this paper, we propose an alternative approach for evaluating and understanding the quality of dMRI data at the subject level using normative modelling. Normative modelling [19] is an innovative method used to model biological and behavioural variation across a study population and can be used to make statistical inferences at an individual level. This is achieved by mapping a response variable (e.g., neuroimaging derived phenotypes) to a covariate (e.g., age) in a similar way growth charts are used in paediatric medicine to map the height or weight of children to their age. Here we demonstrate that using our approach detects subjects with either poor data quality and/or with processing problems as extreme outliers from a normative model that captures population variation within each image derived phenotype (IDP). Crucially: (i) this does not require us to label artefacts in advance, nor (ii) specify what the consequences of different types of artefacts are on the derived phenotype, and (iii) allows immediate identification of potentially problematic scans from large datasets without labour-intensive manual screening. For this purpose, we use the UK Biobank dataset [20,21] which is one of the largest biomedical databases and research resources (currently) containing over 60,000 subjects with available diffusion data as well as hundreds of diffusion IDPs. In a subset of 500 participants we also visually QC and extract quantitative QC parameters to compare with our normative modelling approach.

## 2. Methods

### 2.1 Dataset

We used a subset of the UK Biobank dataset containing 23158 subjects with available dMRI data, from the 2020 data release. Briefly, data were acquired using a Siemens Skyra 3T and had an acquisition time of 7 minutes. It contains 5 x b0 scans, 50 x b1000 s/mm^2^ and 50 x b2000 s/mm^2^ with gradient timings δ = 21.4 ms, Δ = 45.5 ms. The resolution is 2×2×2 mm^3^ and the matrix size is 104×104×72. In this manuscript, we use image-derived phenotypes derived from the standard UKB processing pipelines. In brief, data were corrected for motion, off-resonance (susceptibility and eddy-current induced) and slice dropout artefacts using the Eddy tool [13,14]. The DTIFIT tool within FSL was used to fit the DTI model to the b=1000 shell to calculate measures such as FA and MD. Along with the DTI model a NODDI model was also fit to the data, yielding additional parameters such as intra-cellular volume fraction (ICVF, an index of white matter neurite density) and isotropic or free water volume fraction (ISOVF). The two models were used to extract 675 diffusion IDPs over 75 different white matter tract regions, obtained from skeletonised ROIs of the JUH white matter MNI atlas. More details on data acquisition and pre-processing can be found in the UKB documentation [22] as well as other papers which address the pre-processing and processing pipelines [23,24].

Our methodological pipeline consisted of 2 parallel streams, summarised in Figure 1. The first of these was following the normative modelling framework applied to the whole available UKB dataset (Figure 1A). The second stream involved visual QC and the extraction of quantitative QC measures on a subset of 500 participants (Figure 1B). These measures were then compared to each other as well as to the normative models.

**Figure 1.**
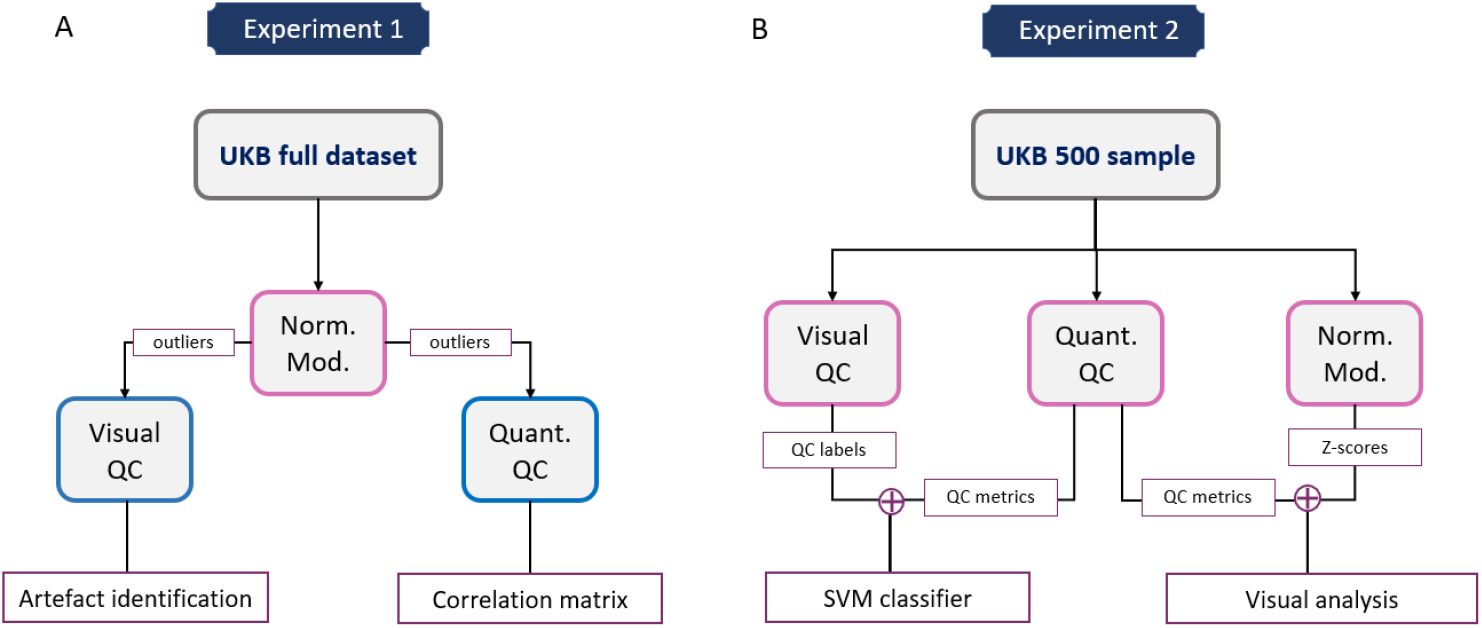
Flowchart describing the main methods. (A) 24 diffusion IDPs provided by UK Biobank were used to fit normative models (norm. mod.). The estimates provided by the models were used in conjunction with the visual QC protocol to identify and label artefact presence in the scans. (B) A subsample of 500 scans were additionally visually rated and had quantitative QC metrics extracted. SVM – support vector machine, QC – quality control, UKB – UK biobank, Norm Mod -Normative model.

### 2.2 Experiment 1

#### 2.2.1 Normative modelling

In preparation for the modelling stage, the subjects with available dMRI were split into test and training sets. The test set consisted of a random sample of 5000 subjects, while the training set consisted of the remainder of the data (n=22658). The list of diffusion IDPs was selected on the basis of diffusion model (DTI and NODDI) and tract, both categories including simple and complex examples with different modelling difficulties. We chose to model across 4 diffusion parameters. The FA and MD parameters of the DTI model were selected for their relative simplicity since they represent a direct measurement of diffusion influenced by tissue microstructure and are the most widely used DTI parameters. We selected the ICVF and ISOVF parameters of the NODDI model for their relative complexity since they enable a more specific characterization of tissue microstructure by estimating neurite density and orientation dispersion. A total of 6 tracts were chosen based on their size and geometrical complexity and comprised: the corpus callosum, the corticospinal tract (both left and right), the uncinate fasciculus (both left and right) and the fornix. This yielded a list of 24 IPDs in total.

A normative model was trained on the training set to estimate the normal range of each IDPs value according to age. To account for the possible non-linear effects and non-Gaussian distributions within the dataset, we used a warped Bayesian linear regression (BLR) model [25], as also used in prior work [25,26]. Specifically, this involves applying a third order polynomial B-spline basis expansion over age with 5 evenly spaced knots with SinhArcsinh warping function described in more detail in Fraza et al. [25]. Next, the test set was used to estimate each subjects’ deviation from the normal range of each IDP by computing the individual z-score (equation 1). The fit statistics of the model were computed including explained variance, skew, and kurtosis. The models were refitted after the exclusion of the outliers from the dataset (see next section) in order to assess the effect the outliers had on the model fit. The amount of deviation for each subject was visualized by plotting the individual z-scores across the mean and centiles of variation predicted by the model. For a detailed introduction to warped BLR normative modelling please consult the dedicated paper by Fraza et al. [25]. All the statistical analyses were performed in Python version 3.8, using the PCN toolkit [27]. We then used the z-statistics derived from the normative model as the basis for further assessment (equation 1).

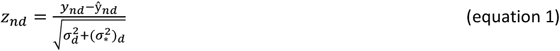

In equation 1, *n* denotes each subject while *d* denotes each IDP and ŷ_*nd*_ is the predicted mean while y_*nd*_ denotes the true response after warping the data to the original input space such that the residuals are as close to Gaussian as possible (see [25] for details). The estimated noise variance (i.e. reflecting variation in the data) is denoted by 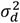 and the variance attributable to modelling uncertainty for the *d*-th IDP is denoted by 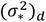

### 2.2.2 Normative modelling outliers

We defined outliers of the normative models as participants who presented a z-score more than 7 standard deviations from the mean in any of the IDPs model. The outlier images were visually assessed and labelled according to the artefact type present. For an accurate label the following scans were inspected: b0 scan (to assess the general quality), T1 scan (to assess the anatomical integrity), skeletonized FA map (to assess registration/model estimation), the FA warp from dMRI space to standard space (to assess the registration quality). We also computed the frequency with which one outlier appears across the IDPs as well as the number of artefact types present in each IDP.

### 2.3 Experiment 2

#### 2.3.1 Visual QC

In a subset of 500 randomly selected participants visual quality control was performed. This consisted of assigning each subject a score from 1 to 3 where 1 = no artefact, 2 = slight artefact, 3 = severe artefact. In order to accelerate the visual QC, 9 slices per subject were used for the visual assessment, 3 from each view (axial, coronal, sagittal) of the B0 image with associated T1 mask (in dMRI space) overlaid. Two experienced raters were trained to perform the manual labelling by following a locally developed protocol which established the types and severity of the artefacts and how to provide a score accordingly. The visually captured artefacts included out of field of view scans, signal loss, brain extraction errors and residual susceptibility distortions. The inter-class correlation (ICC3k within the Pingouin library [28] in Python 3.8) metric was then used to assess the agreement between the two raters.

#### 2.3.2 Quantitative QC

The quantitative quality control (QQC) measurements consisted of 21 quality descriptors for diffusion data including 18 parameters obtained with the Eddy QC tool [15], 2 parameters directly available from UKB and one locally derived parameter. This list includes motion parameters (average and relative motion, translation, rotation), number of outliers per slice (for assessing signal dropout), SNR and angular CNR, T1 vs DWI discrepancy and the discrepancy (i.e., registration cost) between the FA image in MNI space and the FMRIB FA template. A full list of the acquired measurements and their source is presented in Table 1 in Appendix 1.

#### 2.3.3 Subject classification

The labels obtained (per subject) with the visual QC were used as ground truth for training a linear SVM classifier for the quantitative QC measurements (table 1). We separated the scores in two classes where scores 1 and 2 were grouped together into a single class representing subjects with acceptable scans (two classes: 0 = acceptable, 1 = not acceptable). Before classification, the data was balanced by subsampling the artifact-free class to prevent biasing the performance results. The data balancing consisted of creating 20 random subsamples of the artefact-free data that match the sample-size of the artefactual data (which varies each round). The classifier was run iteratively 20 times and the classification performance (measured by accuracy, sensitivity, and specificity) obtained using 5-fold cross validation was averaged across the 20 iterations to obtain the final results.

#### 2.3.4 Comparing normative modelling with visual and quantitative QC

In order to assess the image quality of the outliers and determine the severity and the type of artefacts present we applied the same visual and quantitative QC protocol to the outliers identified with normative modelling. Furthermore, we also identified the quantitative QC outliers (threshold of 2 standard deviations from the mean) and again computed the number of times each outlier appeared across all quantitative QC parameters. A correlation matrix was computed using this information which consisted of the following measurements: T1 discrepancy values, FA vs DWI discrepancy, SNR, Normative Modelling (NM) outlier frequency, quantitative QC outlier frequency and visual QC score to compare between quality descriptors, visual evaluation and the involved IDP.

## 3. Results

### 3.1 Experiment 1

#### 3.1.1 Normative modelling

The normative modelling fit was measured in terms of explained variance (EV), skewness and kurtosis. Across the 24 IDPs the models had mean EV of μ_EV_ = 0.092 with standard deviation σ_EV_ = 0.089, mean Skew of μ_Skew_ = 0.433 with standard deviation σ_Skew_ = 0.686, mean Kurtosis of μ_Kurtosis_ = 1.482 with standard deviation σ_Kurtosis_ = 1.886. The difference in fit of the model with and without outliers was not significant (p = 0.33) as tested with a paired t-test of the z-scores before and after outlier exclusion.

#### 3.1.2 Normative modelling outliers

The normative models were fit to the 24 diffusion IDPs and the subjects with a z-score beyond 7 standard deviations away from the mean were considered outliers. (Figure 2) Across all IDPs there were a total of 85 unique outlier subjects, out of which 64 appeared in at least two IDPs. After visually inspecting each outlier, we devised 3 main categories of image errors (see Figure 3 for examples of each):

**Figure 2.**
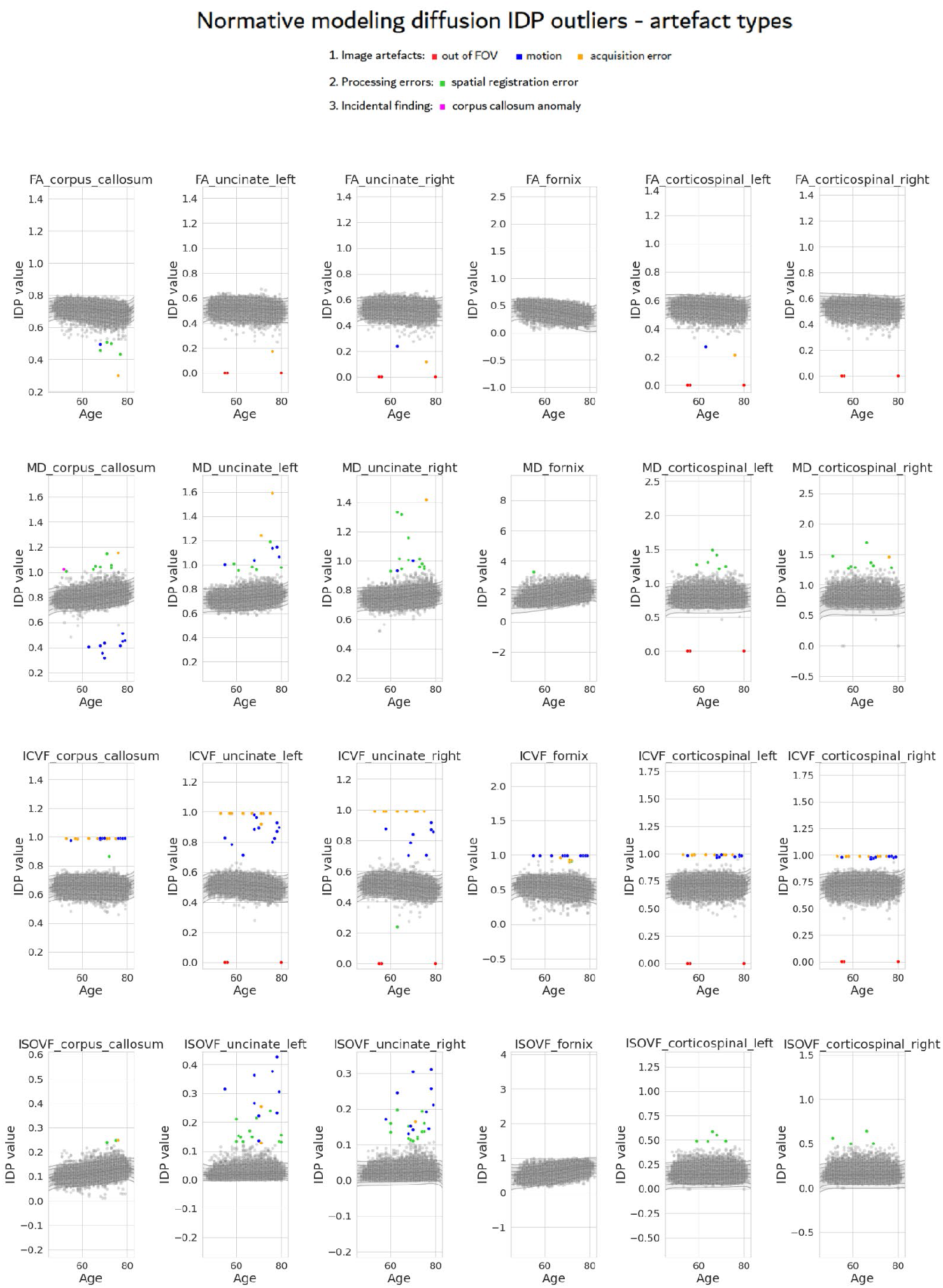
Normative model plots with mean and centiles for each of the 24 diffusion IDPs. Test/Training data is depicted with grey dots. The outliers are colour coded by artefact type, to highlight their position and frequency across different IDPs. Outliers sometimes cluster according to artefact type while also varying depending on DWI parameter and tract.

**Figure 3.**
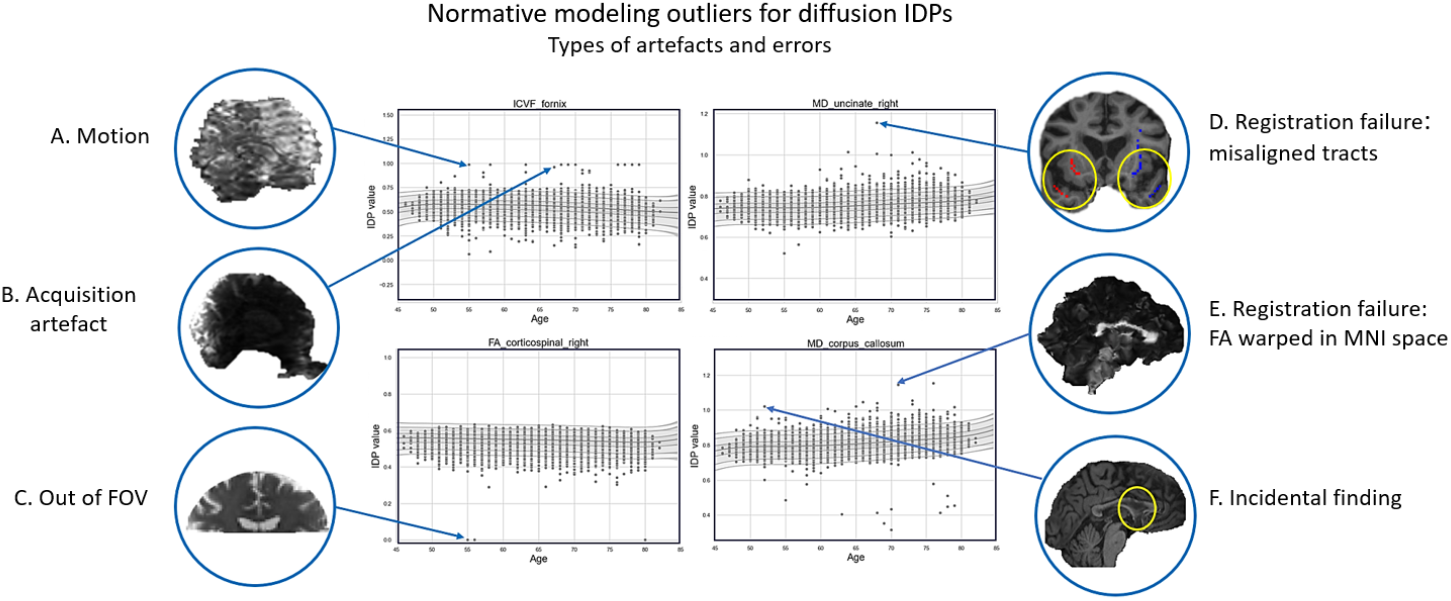
Example of each artefact and error type up close. On the left side 3 A, 3 B and 3 C are examples of image artefacts. On the right side, 3 D and 3 E presents examples of scans with processing errors. 3 F is an example of an incidental finding which was detected as an outlier by the normative model.

1. **Acquisition artefacts**: out of the field of view (out of FOV images i.e., incomplete brain coverage), residual susceptibility or Eddy current artefacts, motion-induced signal dropout etc.
2. **2. Processing errors**: this is mainly comprised of registration or brain extraction errors which can occur for various reasons. Importantly, these errors often arise even from good quality data, e.g., in the case of misregistration due to large ventricles, severe atrophy or anatomical anomalies.
3. **3. Incidental findings**: Although this is not an artefact or error, it is important to identify anatomical anomalies and have the possibility to review such participants to decide whether they should be excluded or not from further analyses.

In Figure 2, we see that the FA metric is robust, showing very few outliers. On the other hand, the ICVF metric across all tracts is more sensitive to image errors, showing many outliers and a high amount of artefact types. Generally, larger tracts such as the corpus callosum and the corticospinal tract are easier to model and have relatively fewer outliers than smaller tracts. The only exception is the fornix, which –perhaps surprisingly– has almost no outliers (except the ICVF), this is because, when looking at the measurement range of the fornix tract IDPs, we can point out the very large variance. This means that the fornix, being a small and hard to model tract is often misregistered which makes this phenomenon not as evident as in the others. When looking at artefact types, it is worth pointing out that MD and ISOVF appear to be more sensitive to processing errors.

Registration failures were uncovered by examining various image types. Figure 3 shows snapshots of examples of different artefacts detected as outliers in the normative model. Figure 3A-C are examples of image artefacts, while Figure 3D is an example of a processing error and specifically a registration failure in a subject with enlarged ventricles who appears as an outlier in the ‘Mean MD in uncinate fasciculus right’, ‘Mean MD in uncinate fasciculus left’ and ‘ISOVF in uncinate fasciculus right’ IDPs. This example is also illustrated in Figure 4, along with the b0 and FA images. The error can be seen in Figure 4C from the overlay of the skeletonized uncinate fasciculus tract from the JHU atlas (in DWI native space) on the T1 scan of the subject (in DWI native space). It is noticeable from the image that the uncinate fasciculus is positioned incorrectly. This indicates a faulty registration and explains why this subject has an abnormal value within the three IDPs mentioned above. Notably, this was not an outlier in other metrics within the same tract despite the mis-registration effecting all downstream images.

**Figure 4.**
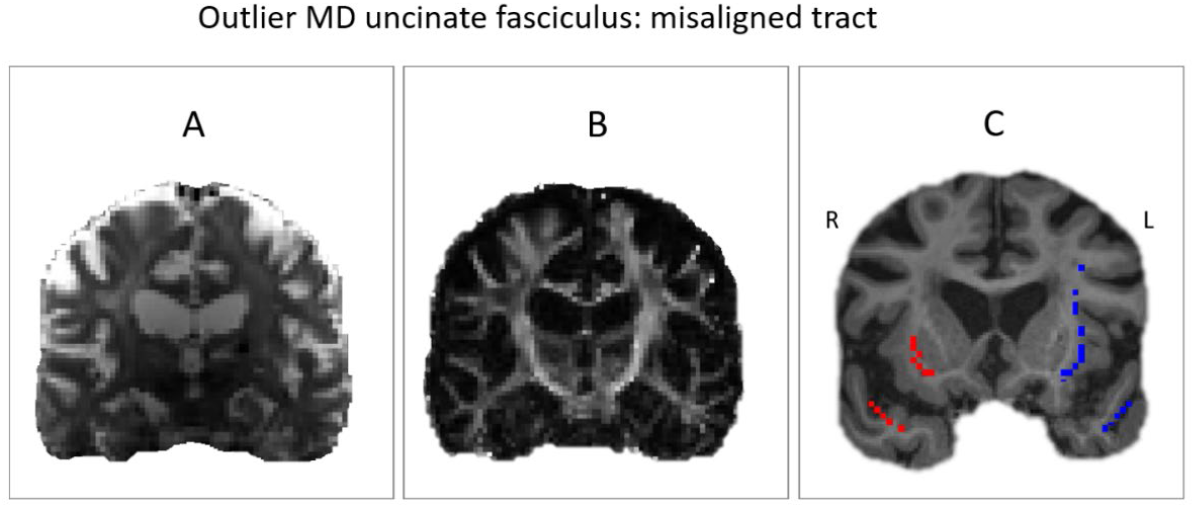
Axial slice of the same subject represented by three different data types. Panel A shows the average b0 image. Panel B shows the FA image while Panel C shows the skeletonized uncinate fasciculus (right: red colour, left = blue colour) overlayed on the T1 image. Close inspection of the registration reveals that the uncinate fasciculus tract is incorrectly aligned with the true white matter tract.

Another example of a processing error in the form of a faulty registration is shown in Figure 3 E which consists of the FA scan warped in MNI space of a subject who appears to be an outlier in the ‘mean MD in corpus callosum’ IDP. It is apparent from the image that the registration is extremely distorted which has a big impact on the appearance of the corpus callosum, thus explaining the abnormal value of this IDP in this subject. Once again it is noteworthy that this participant was not an outlier in other metrics or tracts despite the drastically bad distortion of the image during registration. It is also important to note that these two examples could not be identified as artefactual by visually inspecting any of the DWI scans from the visual protocol. The last example in Figure 3 F shows the T1 scan of a subject with an abnormal corpus callosum which has been identified as an outlier in the ‘mean MD in corpus callosum’. This anatomical abnormality is an incidental finding.

### 3.2 Experiment 2

#### 3.2.1 Visual QC and quantitative QC

Visual QC was performed on a sample of 500 subjects by two raters who gave scores according to the artefact severity from 1 to 3. The raters had an ICC3k score [28] of 0.71 which indicates a moderate reliability and underscores the difficulty in obtaining reliable estimates from manual labelling. In Figure 5.A, SNR values are colour coded in the normative model centile plot of one of the IDPs (mean MD in uncinate fasciculus left). There appears to be little to no correlation between the quantitative QC outliers and the normative modelling outliers but Figure 5A is provided as an example. The visual QC scores were used in several analyses to assess the correlation between visual and quantitative QC measures. In Figure 5B, the visual QC scores of rater 1 were colour coded in the normative model centile plot of one of the IDPs (mean MD in uncinate fasciculus left). There also appears to be little correlation between the unacceptable scans (score = 3) and the normative modelling outliers (data points further away from the mean). Figure 5C also shows a plot of the pairwise relationships between several quantitative QC parameters (T1 discrepancy, SNR and CNR b=1000) and the visual QC scores which are colour coded in the scatterplots. The distributions of the scores overlap consistently and the scatterplots do not show any clear clustering of the data points by score. This suggests that the correlation between the visual and automated QC is weak.

**Figure 5.**
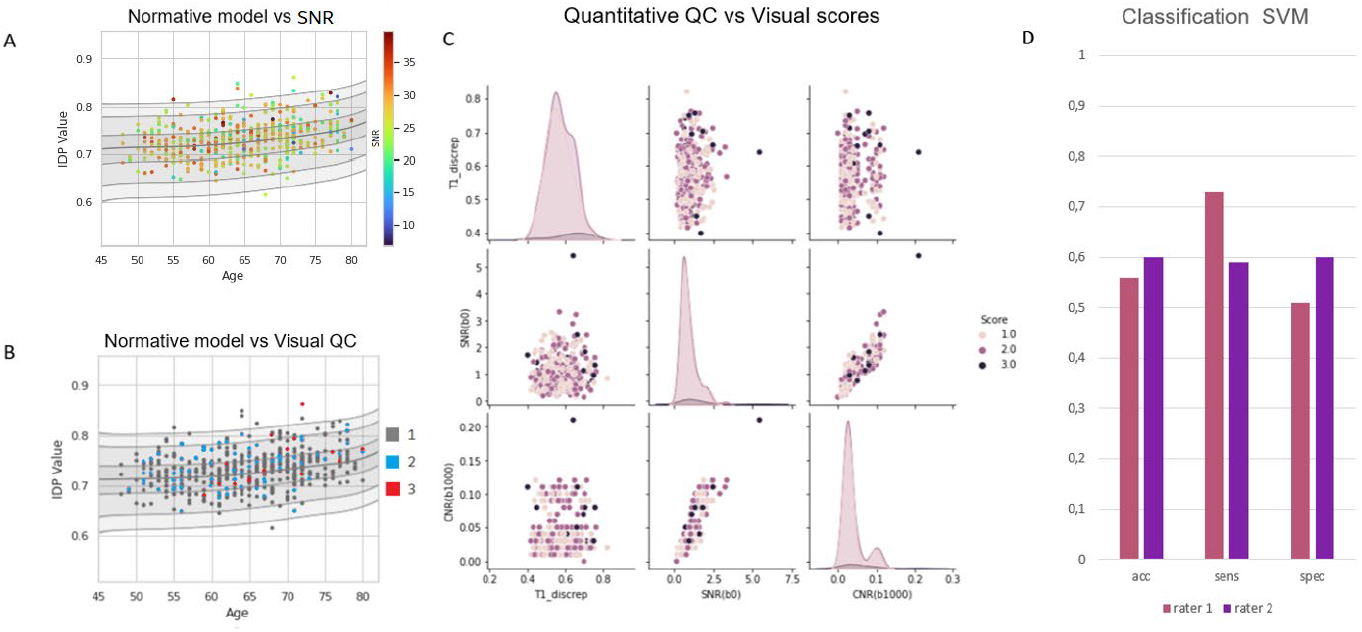
A) SNR values of the 500 subjects colour coded into the normative model centile plot of the ‘mean MD in uncinate fasciculus left’ IDP. B) Visual QC scores of the 500 subjects colour coded into the normative model centile plot of the ‘mean MD in uncinate fasciculus left’ IDP. C) Pairplot between three quantitative QC parameters and the visual QC scores showing the pairwise relationship between visual and automate QC modalities. D. SVM classification performance scores for both raters.

#### 3.2.2 Subject classification

Figure 5 D shows the performance scores of the SVM classifier which was used to distinguish between acceptable (score 1 and 2) and unacceptable scans (score 3). The accuracy scores are no higher than 0.6 in both rates which corroborates that the link between visual and automated QC is not significant.

#### 3.2.3 Comparing normative modelling with visual and quantitative QC

Using the 85 outlier subjects from the larger dataset, a correlation matrix was computed with select measurements obtained from both the visual QC and quantitative QC parameters as well as parameters such as frequency with which a subject appears as outlier in the 24 IDP normative models. This can be seen in Figure 6. In this case there is a strong positive correlation between the visual QC scores and the T1 discrepancy parameter as well as the outlier frequency in both the normative model and the quantitative QC. This suggests that, in the case of extreme outliers, the correlation between QC measures is much stronger.

**Figure 6.**
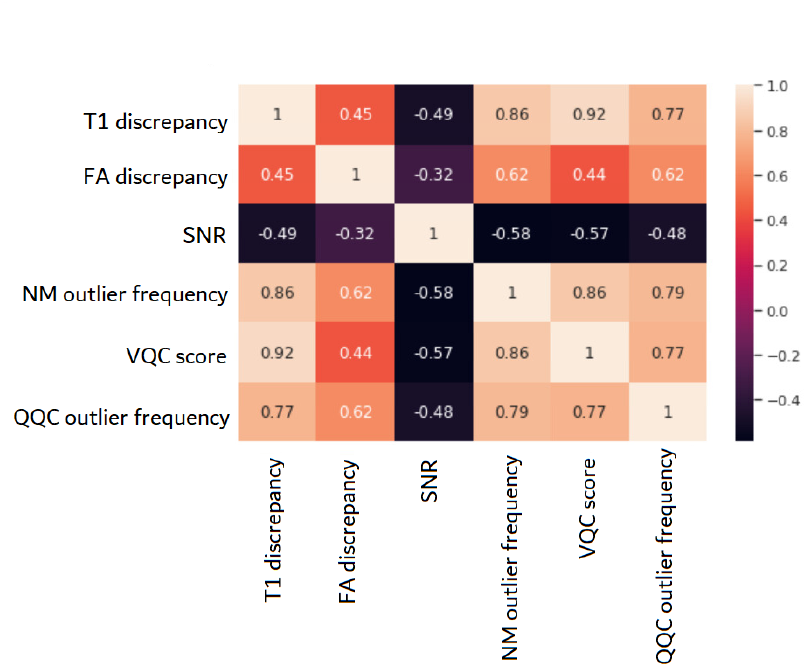
Correlation matrix between quantitative QC (QQC) metrics (T1 discrepancy, FA discrepancy, SNR and QQC outlier frequency), normative modelling (NM) outlier frequency and Visual QC (VQC) score. This correlation was computed using the 85 outliers detected by the normative model and shows the relationship between the three QC methods in case of images heavily affected by artefacts and processing errors.

## 4. Discussion

In this paper we propose a new approach for dMRI quality control. By training normative models for 24 diffusion IDPs and analysing the outliers, we were able to identify severe artefact and processing errors which were not detected by other QC methods such as visual QC or quantitative QC metrics like those obtained with EDDY QC. Therefore, the time and effort invested in assessing the quality of a large dMRI dataset was significantly reduced. We were able to detect processing errors that may occur even in the absence of artefacts and that may go undetected in conventional QC workflows. At the same time, we showed the wide discrepancy between QC approaches and demonstrated that visual QC is subjective and prone to failure since artefacts can be subtle or at times impossible to detect visually.

Another important finding from this work was that we show that artefact detection is dependent on both the type of derived measure and the tract from which it is derived (i.e., regional specificity).

The normative modelling approach to quality control has the advantage of being able to detect processing errors and can be applied easily to any IDP of interest. This is an important achievement due to the high impact these errors can have on the results of an analysis but also as some of the detected errors may be resolvable with re-processing. Conversely, the scans which contain severe image artefacts can often not be corrected and must be discarded. As discussed in the introduction there are several toolboxes dedicated to correcting image artefacts such as susceptibility, motion and eddy induced distortions. However, these algorithms, although robust, can leave behind residual image artefacts which in some cases may render the entire subjects’ scan unusable. Unlike these image artefacts, the processing errors may be resolvable by adjusting the pipelines used and potentially allowing the data to still be used.

We performed visual and automated QC for a sample of 500 UK Biobank subjects to assess the compatibility of the methods. Specifically, we wanted to see if the same outliers occur for all methods and evaluate the amount of overlap. We show that there is very little (visual) overlap between the outliers of the three methods (visual, quantitative, and normative modelling). The performance of the SVM classifier for both raters also suggests a weak link between the manual labels and quantitative parameters and underscores the difficulty in performing manual QC for diffusion data, even for experienced raters. However, the correlation matrix in Figure 6, which was computed on the 85 outliers, tells us that in extreme cases, the three modalities have a close relationship and can explain each other’s variance. We would like to suggest that these results do not necessarily mean that the three methods are incompatible in the case of less severe artefacts but that that are actually complementary. A complete and detailed quality assessment of diffusion MRI scans (if necessary) would make use of all these methods which are designed to detect different artefact types. It would be fair to say, however, that the normative modelling approach to QC is a comprehensive method which can identify a broad range of artefacts, processing errors and incidental findings and can be adapted easily to various needs depending on the scope and aims of the study. The code used to prepare the data and train the models is available online in order [29] to make this method accessible to the public.

A strength of the normative modelling approach is that it does not need any manually labelled data. Many QC methods which are based on classification require accurate ground truth data which must be acquired through a visual QC protocol. We have shown in this study that inter-rater reliability is not perfect, which although seems to be a limitation, only reflects the reality of manual labelling – it is imperfect and subjective. Therefore, carrying out a visual QC protocol is undesirable and makes the classification method unlikely to be preferred. Classification based methods also have the shortcoming of being unable to identify artefacts which are not present in the training set. This is not the case for the normative modelling method, which is able to detect unseen errors. Furthermore, our approach does not suffer from the shortcomings of unbalanced datasets which present a problem in any method which uses artefact vs artefact-free classes. The nature of the dMRI dataset (and most datasets in the medical imaging field) is that most of the scans will be artefact-free, creating a very large imbalance between classes which must be carefully handled to overcome bias in classification performance scores. In this paper we have seen the low frequency with which outliers were observed in the UKB sample, which poses a major problem for manual QC approaches (particularly for large samples) and will only get worse as image processing techniques for diffusion data improve because the artefactual scans will be fewer and more difficult to recognize. The normative modelling approach can assess the level of abnormality at the individual level rendering categorical class labels unnecessary. This is a powerful point since class division puts limitations on interindividual variability and suppresses the dimensionality of the data.

The YTTRIUM method [16] is similar to the normative approach because it uses a measure of distance from the mean to establish an outlying behavior of diffusion QC measurements at the subject level. Although, this method is comprehensive and straight forward, the normative modelling method has the added advantage of customizing the region of interest by virtue of the IDPs which are selected for training the model as well as the choosing the relevant diffusion parameter depending on the study. The normative modelling in the present paper has an emphasis on age which was used as a covariate for training. By selecting age as our covariate, we exposed the increased outlier frequency proportional to age as well as the normal trajectory of each IDP as a function of age (within the used age range i.e., 45-85). However, the normative models can also include other covariates depending on the scope of the study (e.g., sex, site, ventricle size, fluid intelligence etc.). Therefore, the normative modelling approach to QC of diffusion data can be extremely versatile and informative. Furthermore, this quality control approach can also be used for other neuroimaging techniques since it is not limited only to diffusion MRI.

There is no gold standard when it comes to dMRI QC and the best method will depend on the analysis to be conducted. For example, detecting group differences via a univariate group analysis probably has a lower sensitivity to artefacts than a biomarker detection task using a classifier. The perfect QC depends on the goal of the study and the type of the data as well as on the size of the dataset and the time available for QC.

Some shortcomings of our method regard the limited number of chosen IDPs. The QC of the dataset could have been more detailed and accurate if more IDPs were used in the analysis and if the outlier threshold was lower, allowing us to catch the less severe artefacts. However, for the purposes of showing the efficacy of the method, we limited ourselves to a representative number of IDPs and a high threshold to detect the most obvious outliers. Nonetheless, this method is not guaranteed to catch all artefacts. If an artefact can be sufficiently corrected that it lies in the bulk of the normative distribution, then it will not be detected as an outlier unless the threshold for deviation from the normal is lowered. This phenomenon is exemplified in Figure 7 (in Supplement 2), where a scan from a subject with a visual score of 3 (by both raters) is visualized in FSL to reveal a substantial artefact caused by signal loss. In the centile plots belonging to the normative models, it is visible that the subject is, indeed, an outlier in two of the IDP, only if the threshold had been slightly lower. Therefore, the threshold is relevant to the detection of the artefactual subject as much as targeted tract and dMRI measurement (e.g., FA, MD, etc.). The risk of lowering the threshold too much constitutes the inclusion of relevant biological effects in the outlier pool. This is because the periphery of the normative modelling consists of a mix of relevant biological effects and artefacts and separating these two can be very difficult.

In conclusion, this paper introduces a new method for dMRI quality control. The normative modelling approach to QC is a semi-automated pipeline which is able to detect subjects who present image artifacts as well as other processing errors. The analysis is performed at an individual level, overcoming the shortcomings of class division as well as having the ability to detect new, unseen artefacts while not requiring any manually labelled data. Moreover, in this study we showed, with the help of our method, that there are three main categories of image errors: image artifacts which can be detected visually but are irreversible and processing errors which usually go undetected but can potentially be fixed. The normative modelling approach can be used together with other QC methods such as a visual QC protocol or quantitative QC parameters and can also be customized for detection severity (by changing the outlier threshold) based on the scope of the study. Finally, we showed that although QC methods are not consistent between each other nor with the normative modelling method, they do align for severe cases and can be complementary enhancing the efficacy of the overall quality assessment. Normative modelling is therefore a promising tool for semi-automated QC of diffusion data.

## 5. Data and code availability

The data used in the present study is part of the UK Biobank dataset which is available to be downloaded upon completing an access application. More information can be found on the dedicated webpage [30]. The code used to process the data and train the normative models is also available online on GitHub [29].

## 6. Author contributions

Ramona Cirstian: Methodology, Software, Data Curation, Writing -Original Draft

Natalie J. Forde: Conceptualization, Supervision, Methodology, Resources, Data Curation, Writing -Review & Editing

Jesper L.R. Andersson: Writing -Review & Editing

Stamatios N. Sotiropoulos: Supervision, Writing -Review & Editing

Christian F. Beckmann: Supervision, Project administration

Andre F. Marquand: Conceptualization, Supervision, Methodology, Resources: Writing -Review & Editing

## 8. Supplement 1

**Table 1.** List of quantitative dMRI QC metrics obtained using automated QC methods.

| QC metric | Description | Source |
| --- | --- | --- |
| Average absolute motion (mm) | Volume-to-volume motion | EDDY QC |
| Average relative motion (mm) | Volume-to-volume motion | EDDY QC |
| Average x translation (mm) | Volume-to-volume motion | EDDY QC |
| Average y translation (mm) | Volume-to-volume motion | EDDY QC |
| Average z translation (mm) | Volume-to-volume motion | EDDY QC |
| Average x rotation (deg) | Volume-to-volume motion | EDDY QC |
| Average y rotation (deg) | Volume-to-volume motion | EDDY QC |
| Average z rotation (deg) | Volume-to-volume motion | EDDY QC |
| Average stdev x translation (mm) | Within volume motion | EDDY QC |
| Average stdev y translation (mm) | Within volume motion | EDDY QC |
| Average stdev z translation (mm) | Within volume motion | EDDY QC |
| Average stdev x rotation (deg) | Within volume motion | EDDY QC |
| Average stdev y rotation (deg) | Within volume motion | EDDY QC |
| Average stdev z rotation (deg) | Within volume motion | EDDY QC |
| Stdev x axis linear term (Hz/mm) | Eddy current-induced distortion | EDDY QC |
| Stdev y axis linear term (Hz/mm) | Eddy current-induced distortion | EDDY QC |
| Stdev z axis linear term (Hz/mm) | Eddy current-induced distortion | EDDY QC |
| Total number of outliers (%) | Slice drop-out | EDDY QC |
| Volume-wise number of outliers (%) | Slice drop-out | EDDY QC |
| Outliers per b-shell (%) | Slice drop-out | EDDY QC |
| Outliers per PE dir (%) | Slice drop-out | EDDY QC |
| Average b0 SNR | General scan quality | EDDY QC |
| Average diffusion CNR per shell | General scan quality | EDDY QC |
| Number of slices corrected | Slice drop-out | UK Biobank |
| T1 vs dMRI discrepancy | Movement artefacts | UK Biobank |
| FA discrepancy | Movement artefacts | Locally derived |

## 9. Supplement 2

**Figure 7.**
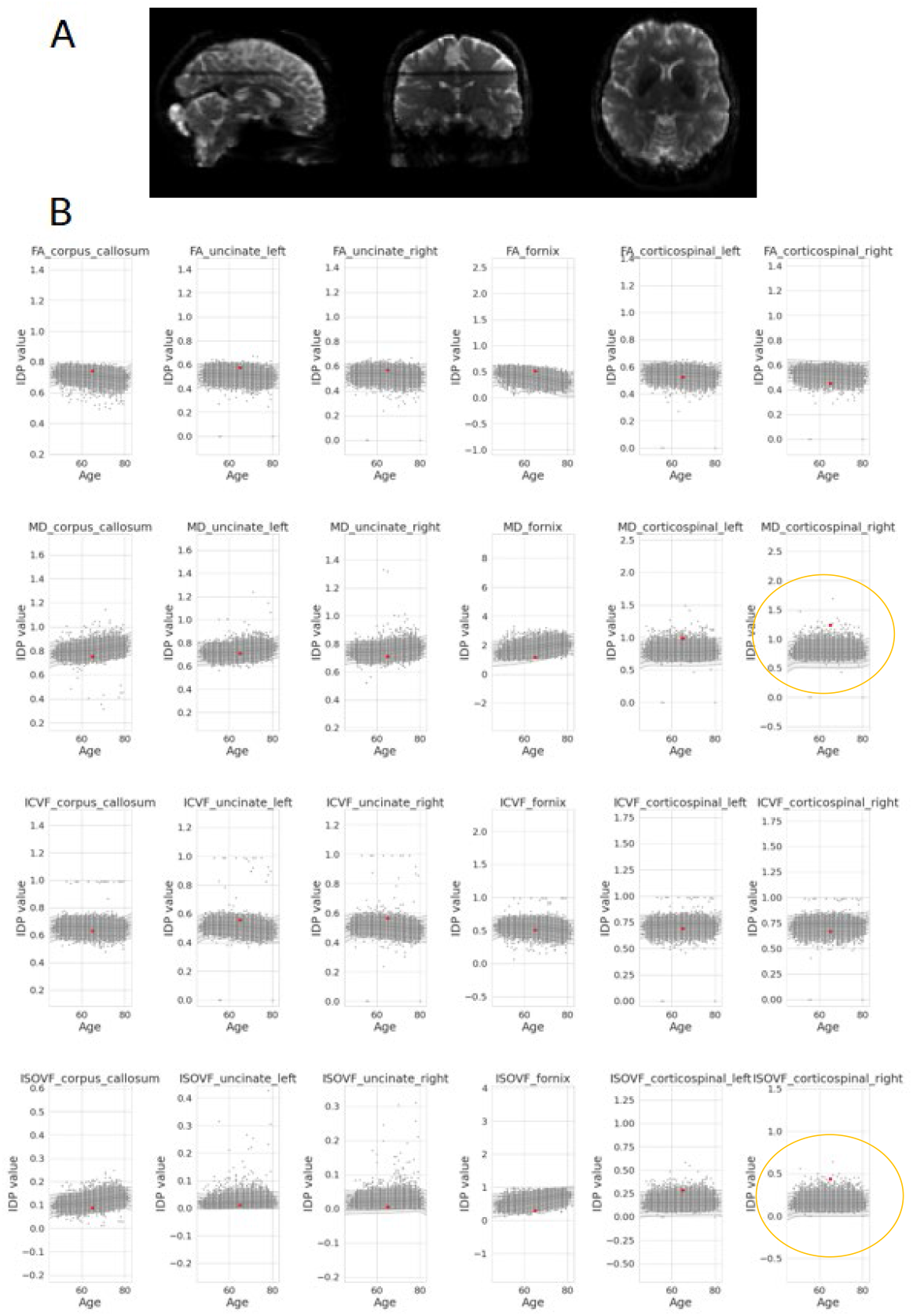
Example of subject (id: 5700767) scored 3 by both raters but unidentified as outliers by normative model. 7A. FSL view of the b0 scan of the subject shows severe signal loss. 7B. The centile plots of the normative models across 24 diffusion IDPs reveal that this subject is an outlier in two IDPs (MD corticospinal tract right and ISOVF corticospinal tract right) if the threshold would be slightly lower than 7 standard deviation from the mean.

## References

1. Le Bihan, D. (2003). Looking into the functional architecture of the brain with diffusion MRI. Nature reviews neuroscience, 4(6), 469–480.

2. Le Bihan, D., Poupon, C., Amadon, A., & Lethimonnier, F. (2006). Artifacts and pitfalls in diffusion MRI. Journal of Magnetic Resonance Imaging: An Official Journal of the International Society for Magnetic Resonance in Medicine, 24(3), 478–488.

3. Le Bihan, D., Mangin, J. F., Poupon, C., Clark, C. A., Pappata, S., Molko, N., & Chabriat, H. (2001). Diffusion tensor imaging: concepts and applications. Journal of Magnetic Resonance Imaging: An Official Journal of the International Society for Magnetic Resonance in Medicine, 13(4), 534–546.

4. Zhang, H., Schneider, T., Wheeler-Kingshott, C. A., & Alexander, D. C. (2012). NODDI: practical in vivo neurite orientation dispersion and density imaging of the human brain. Neuroimage, 61(4), 1000–1016.

5. Andersson, J. L. (2021). Diffusion MRI artifact correction. In Advances in Magnetic Resonance Technology and Applications (Vol. 4, pp. 123–146). Academic Press.

6. Wu, Y. C., Harezlak, J., Elsaid, N. M., Lin, Z., Wen, Q., Mustafi, S. M., … & McAllister, T. W. (2020). Longitudinal white-matter abnormalities in sports-related concussion: A diffusion MRI study. Neurology, 95(7), e781–e792.

7. Lepage, C., de Pierrefeu, A., Koerte, I. K., Coleman, M. J., Pasternak, O., Grant, G., … & Bouix, S. (2018). White matter abnormalities in mild traumatic brain injury with and without posttraumatic stress disorder: a subject-specific diffusion tensor imaging study. Brain imaging and behavior, 12, 870–881.

8. Ho, T. C., Sisk, L. M., Kulla, A., Teresi, G. I., Hansen, M. M., Wu, H., & Gotlib, I. H. (2021). Sex differences in myelin content of white matter tracts in adolescents with depression. Neuropsychopharmacology, 46(13), 2295–2303.

9. Meinert, S., Repple, J., Nenadic, I., Krug, A., Jansen, A., Grotegerd, D., … & Dannlowski, U. (2019). Reduced fractional anisotropy in depressed patients due to childhood maltreatment rather than diagnosis. Neuropsychopharmacology, 44(12), 2065–2072.

10. Chang, L. C., Walker, L., & Pierpaoli, C. (2012). Informed RESTORE: a method for robust estimation of diffusion tensor from low redundancy datasets in the presence of physiological noise artifacts. Magnetic resonance in medicine, 68(5), 1654–1663.

11. Zwiers, M. P. (2010). Patching cardiac and head motion artefacts in diffusion-weighted images. Neuroimage, 53(2), 565–575.

12. Oguz, I., Farzinfar, M., Matsui, J., Budin, F., Liu, Z., Gerig, G., … & Styner, M. (2014). DTIPrep: quality control of diffusion-weighted images. Frontiers in neuroinformatics, 8, 4.

13. Andersson JL, Sotiropoulos SN. An integrated approach to correction for off-resonance effects and subject movement in diffusion MR imaging. Neuroimage. 2016 Jan 15;125:1063-78.

14. Andersson, J. L., & Sotiropoulos, S. N. (2016). An integrated approach to correction for off-resonance effects and subject movement in diffusion MR imaging. Neuroimage, 125, 1063–1078.

15. Bastiani, M., Cottaar, M., Fitzgibbon, S. P., Suri, S., Alfaro-Almagro, F., Sotiropoulos, S. N., … & Andersson, J. L. (2019). Automated quality control for within and between studies diffusion MRI data using a non-parametric framework for movement and distortion correction. Neuroimage, 184, 801–812.

16. Maximov, I. I., van Der Meer, D., de Lange, A. M. G., Kaufmann, T., Shadrin, A., Frei, O., … & Westlye, L. T. (2021). Fast qualitY conTrol meThod foR derIved diffUsion Metrics (YTTRIUM) in big data analysis: UK Biobank 18,608 example. Human brain mapping, 42(10), 3141–3155.

17. Samani, Z. R., Alappatt, J. A., Parker, D., Ismail, A. A. O., & Verma, R. (2020). QC-Automator: Deep learning-based automated quality control for diffusion mr images. Frontiers in neuroscience, 13, 1456.

18. Ahmad, A., Parker, D., Dheer, S., Samani, Z. R., & Verma, R. (2023). 3D-QCNet–A pipeline for automated artifact detection in diffusion MRI images. Computerized Medical Imaging and Graphics, 103, 102151..

19. Marquand, A. F., Rezek, I., Buitelaar, J., & Beckmann, C. F. (2016). Understanding heterogeneity in clinical cohorts using normative models: beyond case-control studies. Biological psychiatry, 80(7), 552–561.

20. Ollier, W., Sprosen, T., & Peakman, T. (2005). UK Biobank: from concept to reality.

21. UK Biobank (2006). Protocol for a large-scale prospective epidemiological resource. www.ukbiobank.ac.uk/resources/.

22. Smith S.M., Alfaro-Almagro F. and Miller K.L. UK Biobank Brain Imaging Documentation version 1.8.

23. Alfaro-Almagro, F., Jenkinson, M., Bangerter, N. K., Andersson, J. L., Griffanti, L., Douaud, G., … & Smith, S. M. (2018). Image processing and Quality Control for the first 10,000 brain imaging datasets from UK Biobank. Neuroimage, 166, 400–424.

24. Miller, K. L., Alfaro-Almagro, F., Bangerter, N. K., Thomas, D. L., Yacoub, E., Xu, J., … & Smith, S. M. (2016). Multimodal population brain imaging in the UK Biobank prospective epidemiological study. Nature neuroscience, 19(11), 1523–1536.

25. Fraza, C. J., Dinga, R., Beckmann, C. F., & Marquand, A. F. (2021). Warped Bayesian linear regression for normative modelling of big data. Neuroimage, 245, 118715.

26. Rutherford, S., Fraza, C., Dinga, R., Kia, S. M., Wolfers, T., Zabihi, M., … & Marquand, A. F. (2022). Charting brain growth and aging at high spatial precision. elife, 11, e72904.

27. https://github.com/amarquand/PCNtoolkit

28. https://pingouin-stats.org/build/html/generated/pingouin.intraclass_corr.html

29. https://github.com/ramonacirstian/Diffusion

30. https://www.ukbiobank.ac.uk/enable-your-research/apply-for-access

